# A chromosome-level genome assembly and annotation of a maize elite breeding line Dan340

**DOI:** 10.1101/2021.04.26.441299

**Authors:** Yikun Zhao, Yuancong Wang, De Ma, Guang Feng, Yongxue Huo, Zhihao Liu, Ling Zhou, Yunlong Zhang, Liwen Xu, Liang Wang, Han Zhao, Jiuran Zhao, Fengge Wang

## Abstract

**Background:** Maize is not only one of the most important crops grown worldwide for food, forage, and biofuel, but also an important model organism for fundamental research in genetics and genomics. Owing to its importance in crop science, genetics and genomics, several reference genomes of common maize inbred line (genetic material) have been released, but some genomes of important genetic germplasm resources in maize breeding research are still lacking. The maize cultivar Dan340 is an excellent backbone inbred line of the Luda Red Cob Group with several desirable characteristics, such as disease resistance, lodging resistance, high combining ability, and wide adaptability.

**Findings:** In this study, we constructed a high-quality chromosome-level reference genome for Dan340 by combining PacBio long HiFi sequencing reads, Illumina short reads and chromosomal conformational capture (Hi-C) sequencing reads. The final assembly of the Dan340 genome was 2,348.72 Mb, including 2,738 contigs and 2,315 scaffolds with N50 of 41.49 Mb and 215.35 Mb, respectively. Repeat sequences accounted for 73.40% of the genome size and 39,733 protein-coding genes were annotated. Analysis of genes in the Dan340 genome, together with those from B73, Mo17 and SK, were clustered into 27,654 gene families. There were 1,806 genes from 359 gene families that were specific to Dan340, of which many had functional gene ontology annotations relating to “porphyrin-containing compound metabolic process”, “tetrapyrrole biosynthetic process”, and “tetrapyrrole metabolic process”.

**Conclusions:** The completeness and continuity of the genome were comparable to those of other important maize inbred lines. The assembly and annotation of this genome not only facilitates our understanding about of intraspecific genome diversity in maize, but also provides a novel resource for maize breeding improvement.

**Research Areas:** Genetics and Genomics; Agriculture, Plant Genetics

**Data Description:** 

## Background

Maize (*Zea mays* ssp. *mays* L., NCBI:txid381124) is one of the most important crops grown worldwide for food, forage, and biofuel, with an annual production of more than 1 billion tons [1]. Owing to rapid human population growth and economic demand, maize has been predicted to account for 45% of total cereal demand by the year 2050 [2]. In addition, it is an important model organism for fundamental research in genetics and genomics [3].

Because of its importance in crop science, genetics and genomics, several reference genomes of common maize inbred lines used in breeding have been released since 2009 [4–8]. However, comparative genomic analyses have found that maize genomes exhibit high levels of genetic diversity among different inbred lines [1, 7, 9].

Meanwhile, accumulating studies have suggested that one or a few reference genomes cannot fully represent the genetic diversity of a species [7, 10, 11].

The maize cultivar Dan340 is an excellent backbone inbred line of the Luda Red Cob Group that has several desirable characteristics, such as disease resistance, lodging resistance, high combining ability, and wide adaptability. More than 50 maize hybrid breeds have been derived from Dan340 since 2000, and their planting area has reached 19 million ha. It is considered that Dan340 originated from a landrace in China and exhibits large genetic differences from other maize germplasms that represent the most important core maize germplasms in China [12]. Therefore, it could serve as a model inbred line for the genetic dissection of desirable agronomic traits, combining ability, heterosis, and breeding history.

In the present study, we constructed a high-quality chromosome-level reference genome for Dan340 by combining PacBio long HiFi sequencing reads, Illumina short reads and chromosomal conformational capture (Hi-C) sequencing reads. The completeness and continuity of the genome were comparable with those of other important maize inbred lines, B73 [4], Mo17 [7], SK [13], PH207 [5], and HZS [8]. Furthermore, comparative genomic analyses were performed between Dan340 and other maize lines, and genes and gene families that were specific to Dan340 were identified. In addition, large numbers of structural variations between Dan340 and other maize inbred lines were detected. The assembly and annotation of this genome will not only facilitate our understanding of intraspecific genomic diversity in maize, but also provides a novel resource for maize breeding improvement.

### Plant materials and DNA sequencing

Inbred line Dan340 (Fig. 1) was selected for genome sequencing and assembly because it is an elite maize cultivar, that plays an important role in maize breeding and genetic research. The plants were grown at 25°C in a greenhouse of the Beijing Academy of Agriculture and Forestry Sciences, Beijing, China. Fresh and tender leaves were harvested from the best-growing individual and immediately frozen in liquid nitrogen, followed by preservation at −80 °C in the laboratory prior to DNA extraction. Genomic DNA was extracted from the leaf tissue of a single plant using the DNAsecure Plant Kit (Tiangen Biotech Co., Ltd., Beijing, China). To ensure that DNA extracts were useable for all types of genomic libraries, the quality and quantity were evaluated using a NanoDrop 2000 spectrophotometer (NanoDrop Technologies, Wilmington, DE, USA) and electrophoresis on a 0.8% agarose gel, respectively.

**Figure 1.**
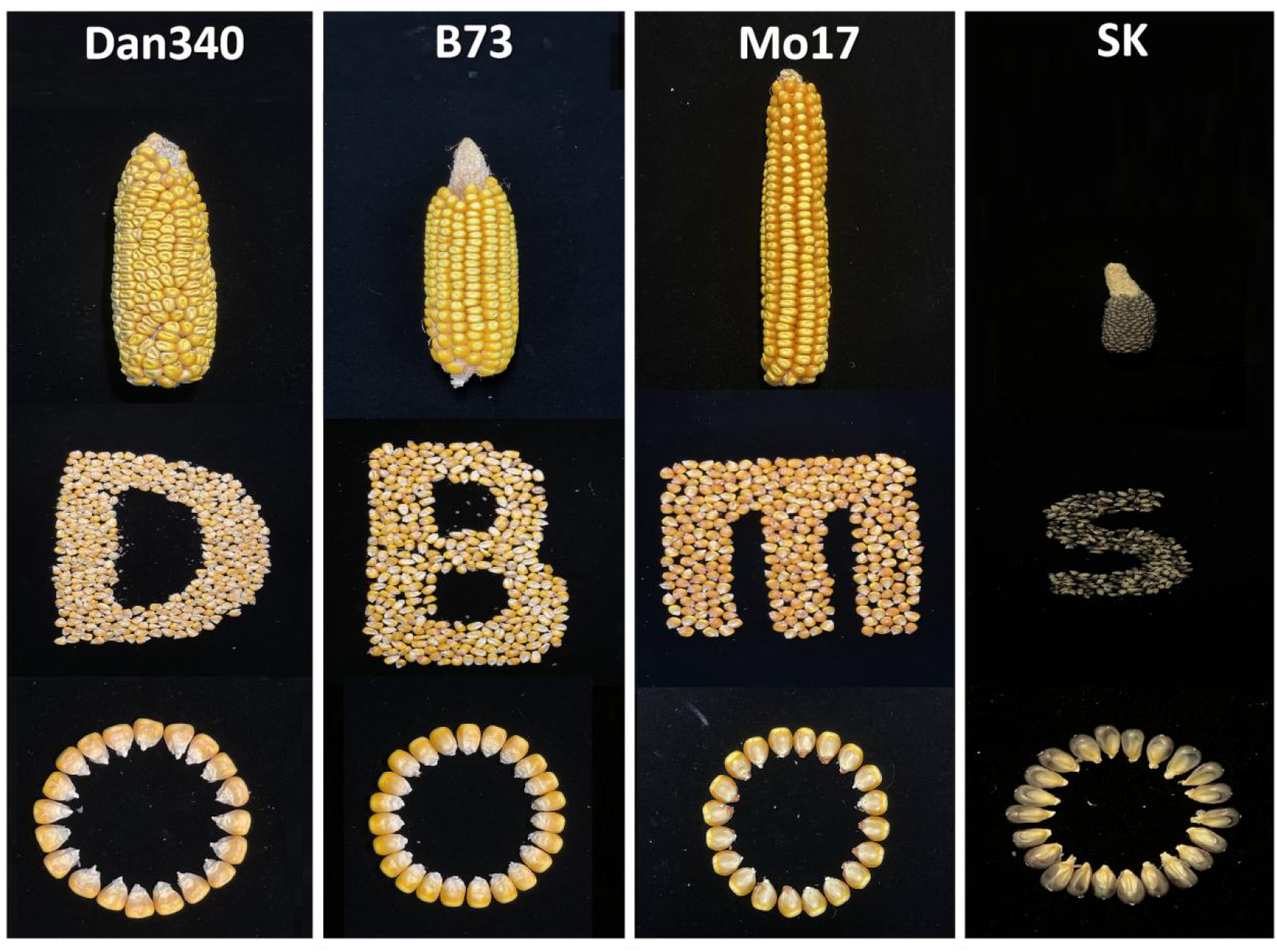
Ear appearances of the maize inbred lines Dan340, B73, Mo17, and SK.

In recent years, third-generation DNA sequencing technologies have undergone rapid technological innovation and are now widely used in genome assembly. In this study, PacBio CCS libraries were prepared using the SMRTbell Express Template Prep Kit 2.0 (Pacific Biosciences, Menlo Park, CA, USA; Ref. No. 101-685-400), following the manufacturer’s protocols, and they were subsequently sequenced on the PacBio sequel II platform (Pacific Biosciences, RRID:SCR_017990). As a result, 63.53 Gb (approximately 27× coverage) of HiFi reads was generated and used for the genome assembly.

In addition, one Illumina paired-end sequencing library, with an insert size of 350 bp, was generated using the NEB Next Ultra DNA Library Prep Kit (NEB, Ipswich, MA, USA) following the manufacturer’s protocol, and it was subsequently sequenced using an Illumina HiSeq X Ten platform (Illumina, San Diego, CA, USA, RRID:SCR_016385) at the Novogene Bioinformatics Institute, Beijing, China. Approximately 80.66 Gb (~34×) of Illumina sequencing data were obtained.

One Hi-C library was constructed using young leaves following previously published procedures, with slight modifications [14]. In brief, approximately 5-g leaf samples from seedling were cut into minute pieces and cross-linked by 4% formaldehyde solution at room temperature in a vacuum for 30 min. Then, each sample was mixed with excess 2.5 M glycine to quench the crosslinking reaction for 5 min and then placed on ice for 15 min. The cross-linked DNA was extracted and then digested for 12 h with 20 units of *DpnII* restriction enzyme (NEB) at 37 °C, and the resuspended mixture was incubated at 62 °C for 20 min to inactivate the restriction enzyme. The sticky ends of the digested fragments were biotinylated and proximity ligated to form enriched ligation junctions and then ultrasonically sheared to a size of 200 - 600 bp. The biotin-labelled DNA fragments were pulled down and ligated with Illumina paired-end adapters, and then amplified by PCR to produce the Hi-C sequencing library. The library was sequenced using an Illumina HiSeq X Ten platform with 2 × 150 bp paired-end reads (Illumina, San Diego, CA, USA). After removing low-quality sequences and trimming adapter sequences, 304.37 Gb (approximately 130×) of clean data were generated and used for the genome assembly.

### Genome assembly

To obtain a high-quality genome assembly of Dan340, we employed both PacBio HiFi reads and Illumina short reads, with scaffolding informed by high-throughput chromosomal conformation capture (Hi-C).

The assembly was performed in a stepwise fashion. First, de novo assembly of the long CCS reads generated from PacBio SMRT sequencing was performed using Hifiasm [15] (RRID:SCR_021069)(https://github.com/chhylp123/hifiasm). A total of two SMRT cells produced 4,073,418 subreads, with an average length of 15,598 bp and a read N50 of 15,715 bp. Generation of HiFi reads and adapter trimming were performed using PacBio SMRTLink v8.0 [16] with default parameters, followed by the deduplication of reads with pbmarkdup v0.2.0 [17] as recommended by PacBio. Next, HiFi reads were aligned to each other and assembled into genomic contigs using Hifiasm [15] with default parameters. The produced primary contigs (p-contigs) were then polished using Quiver [18] by aligning SMRT reads. Then, Pilon [19] (RRID:SCR_014731) was used to perform the second round of error correction with short paired-end reads generated from Illumina Hiseq platforms. Subsequently, the Purge Haplotigs pipeline [20] was used to remove redundant sequences formed as a result of heterozygosity. The draft genome assembly was 2348.68 Mb, which reached a high level of continuity, with a contig N50 length of 45.11 Mb.

For Hi-C reads, to avoid reads having an artificial bias, we removed the following type of reads using HICUP software [21] (RRID:SCR_005569)(http://www.bioinformatics.babraham.ac.uk/projects/hicup/): (a) Reads with ≥ 10% unidentified nucleotides (N); (b) Reads with > 10 nt aligned to the adapter, allowing ≤ 10% mismatches; and (c) Reads with > 50% bases having a phred quality < 5. The filtered Hi-C reads were aligned against the contig assemblies using BWA (version 0.7.8, RRID:SCR_010910)(http://bio-bwa.sourceforge.net/). Reads were excluded from subsequent analyses if they did not align within 500 bp of a restriction site or did not uniquely map, and the number of Hi-C read pairs linking each pair of scaffolds was tabulated. ALLHiC v0.8.12 [22] was used in simple diploid mode to scaffold the genome and optimize the ordering and orientation of each clustered group, producing a chromosome-level assembly. Juicebox Assembly Tools v1.9.8 [23] (RRID:SCR_021172) was used to visualize and manually correct the large-scale inversions and translocations to obtain the final pseudo-chromosomes (Fig. 2). Finally, a total of 2315 scaffolds (representing 91.30% of the total length) were anchored to 10 chromosomes (Fig. 3).

**Figure 2:**
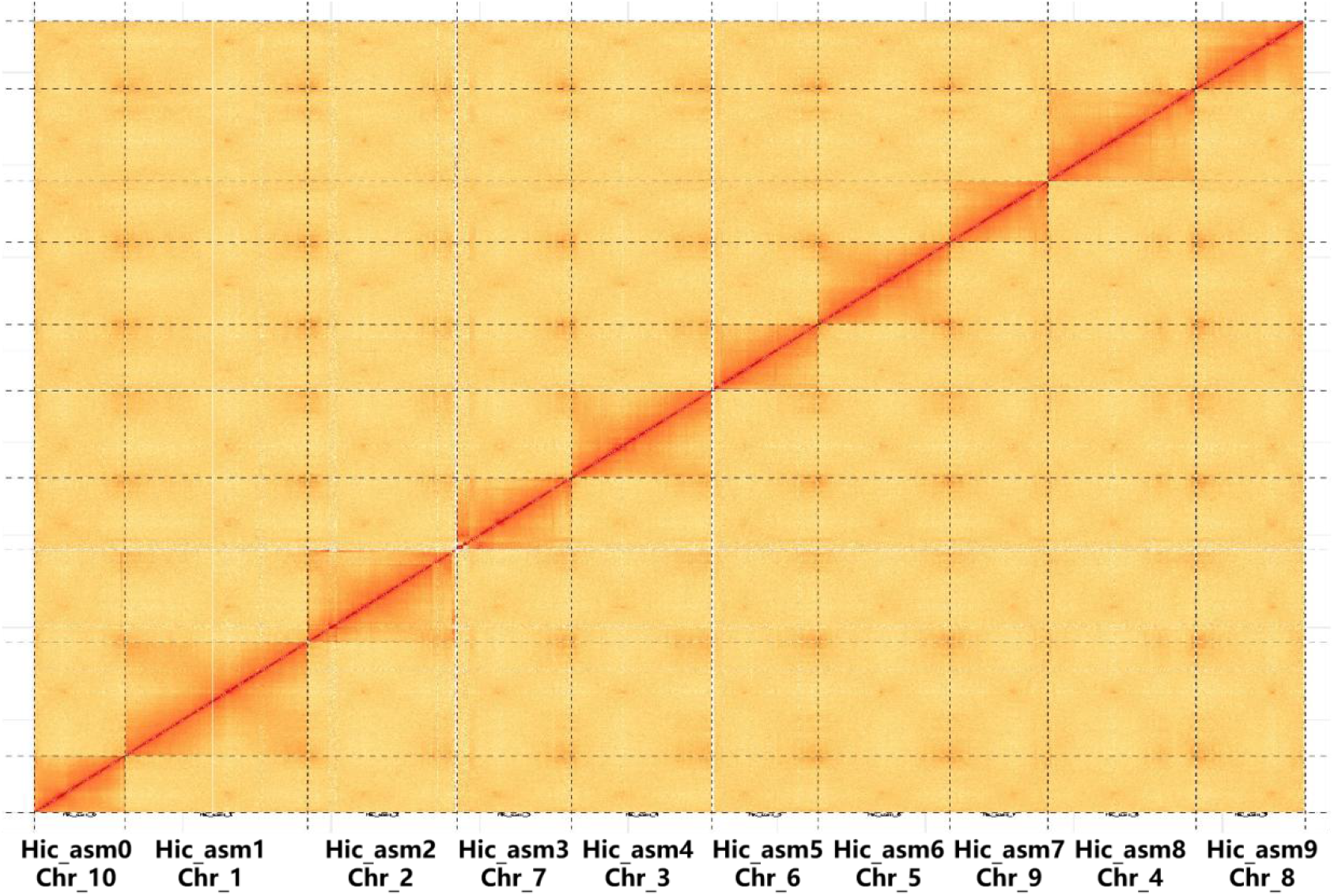
Hi-C contact heat map displaying the inter- and intra-chromosomal interactions in maize inbred line Dan340 genome.

**Figure 3.**
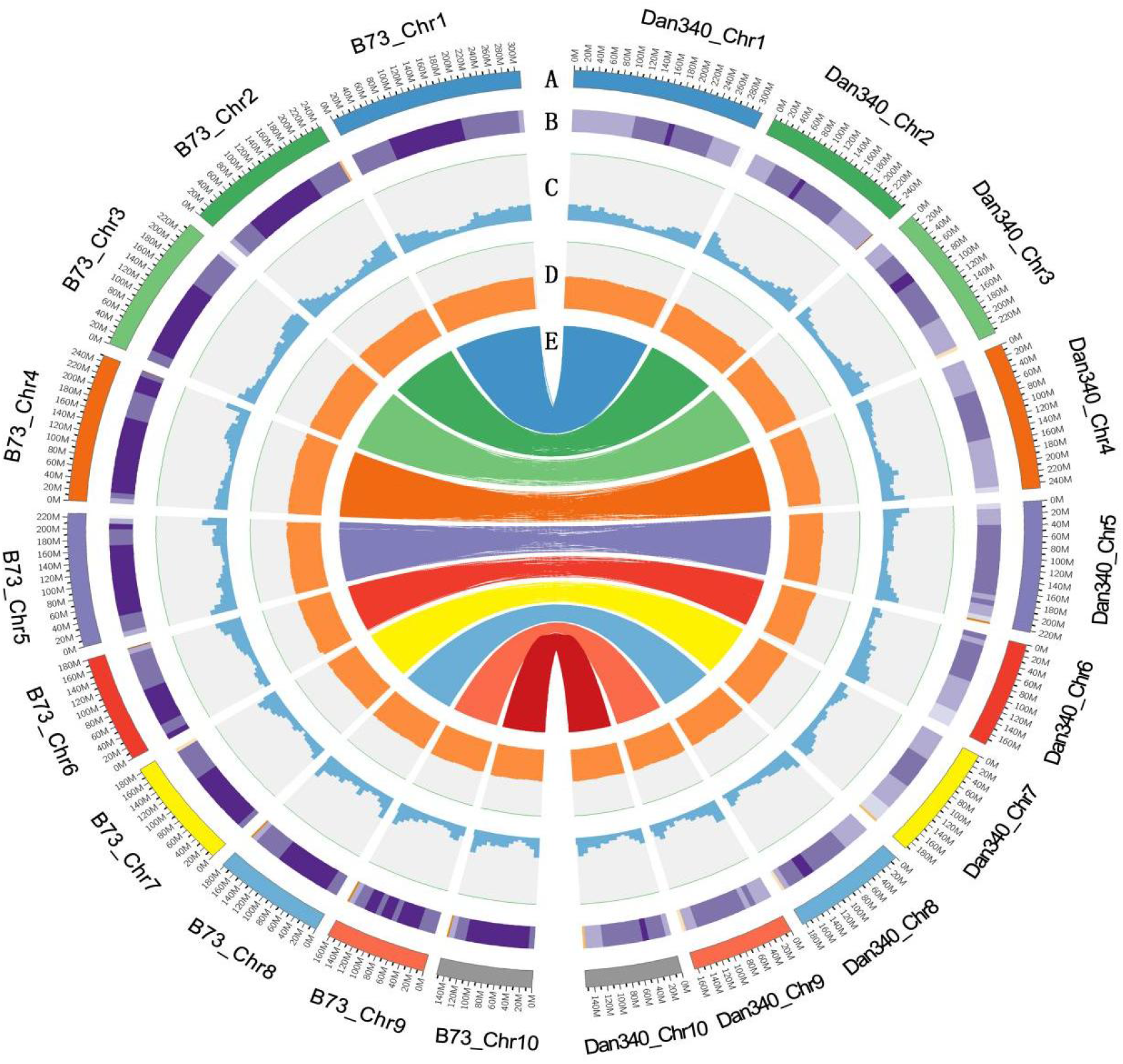
Circos plot of genomic features. Outer-to-inner tracks indicate the following: A, Chromosome numbers of Dan340 and B73; B, Repeat density; C, Histogram of gene density distributions along the chromosomes; D, Histogram of GC content distributions along the chromosomes; E, Syntenic relationships of gene pairs between Dan340 and B73 genomes identified using the best-hit method.

The final assembly of the Dan340 genome was 2,348.72 Mb, including 2,738 contigs and 2,315 scaffolds, with N50 of 41.49 Mb and 215.35 Mb, respectively (Table 1).

**Table 1.** Genome assembly and annotation statistics for the four tested maize inbred lines.

| Genomic features | Dan340 | B73 | Mo17 | SK |
| --- | --- | --- | --- | --- |
| assembled genome size (bp) | 2,348,678,871 | 2,182,075,994 | 2,104,465,715 | 2,161,392,594 |
| Number of scaffolds | 2315 | 687 | 2203 | 671 |
| Total length of scaffolds (Mb) | 2,348.72 | 2,182 | 2,182 | 2,162 |
| Scaffold N50 | 222,765,871 | 226,353,449 | 220,382,597 | 73,237,962 |
| Number of contigs | 2,738 | 1,395 | 9,040 | 1,090 |
| Total length of contigs (Mb) | 2,144,444 | 2,178,268 | 2,147,495 | 2,150,874 |
| Contig N50 | 45,109,016 | 47,037,903 | 1,491,782 | 15,776,512 |
| Number of genes | 39,733 | 39,756 | 38,620 | 43,271 |

### Evaluation of assembly quality

We assessed the quality of the assembly using several independent methods. First, the short reads obtained from the Illumina sequencing data were aligned to the final assembly using BWA version 0.7.8 [24]. Our result showed that the percent of reads mapped to the reference genome was up to 97.48%. Second, a total of 248 conservative genes existing in six eukaryotic model organisms were selected to form the core gene library for the CEGMA (Core Eukaryotic Genes Mapping Approach) [25] (RRID:SCR_015055) evaluation. Our assembled Dan340 genome was aligned to this core gene library using TBLASTN [26] (RRID:SCR_011822), Genewise version 2.2.0 [27] (RRID:SCR_015054), and Geneid v1.4 [28] (RRID:SCR_021639) software tools to evaluate its integrity. The result showed that 238 complete (95.97%) and 243 partial (97.98%) genes were detected in our assembly. Third, the completeness was assessed using the benchmarking universal single-copy orthologs (BUSCO) [29] (RRID:SCR_015008). The final assembly was tested against the database embryophyta_odb10, which includes 1,614 conserved core genes. The result showed that 98.08% (1,583), 1.11 % (18), and 0.81% (13) of the plant single-copy orthologs were present in the assembled Dan340 genome as complete, fragmented, and missing genes, respectively. Fourth, the long-terminal repeat (LTR) Assembly Index (LAI) metric was used to evaluate assembly continuity in Dan340 and three other maize genomes (B73, Mo17 and SK, Fig. 4). Intact LTR retrotransposons in the four genomes were identified using LTRharvest v1.6.1 [30] (RRID:SCR_018970), LTR_FINDER v1.07 [31] (RRID:SCR_015247), and LTR_retriever v2.9.0 [32] (RRID:SCR_017623). The LAI pipeline was executed using the following parameter settings: -t 20 -intact genome.fasta.pass.list -all genome.ltr.fasta.out. Our Dan340 genome had a LAI score of 25.13, which was relatively high among the four maize genomes compared in this study. B73, Mo17, and SK produced scores of 24.94, 24.45, and 27.12, respectively (Fig. 4 and Table 2). A higher LAI score indicates more complete genome assembly because more intact LTR retrotransposons are identified, as in our Dan340 genome. Furthermore, whole-genome sequence alignments of Dan340 to the genomes of the other three maize inbred lines demonstrated that our assembly has highly collinear relationships with other published maize genomes (Fig. 5). Taken together, the assessment results suggested that the Dan340 genome assembly was of high quality.

**Figure 4.**
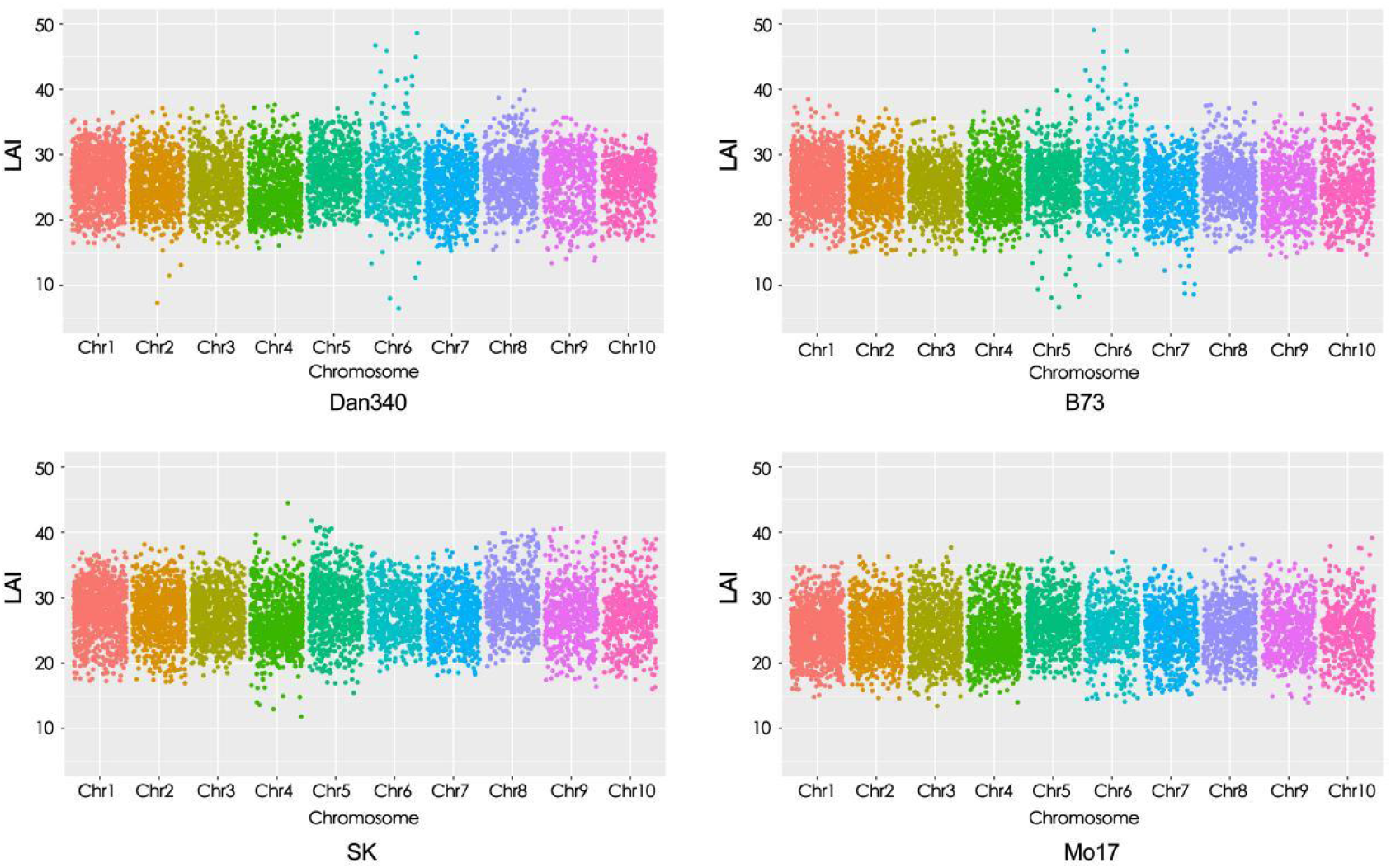
Genome-wide LTR Assembly Index (LAI) scores for Dan340, B73, Mo17 and SK.

**Figure 5.**
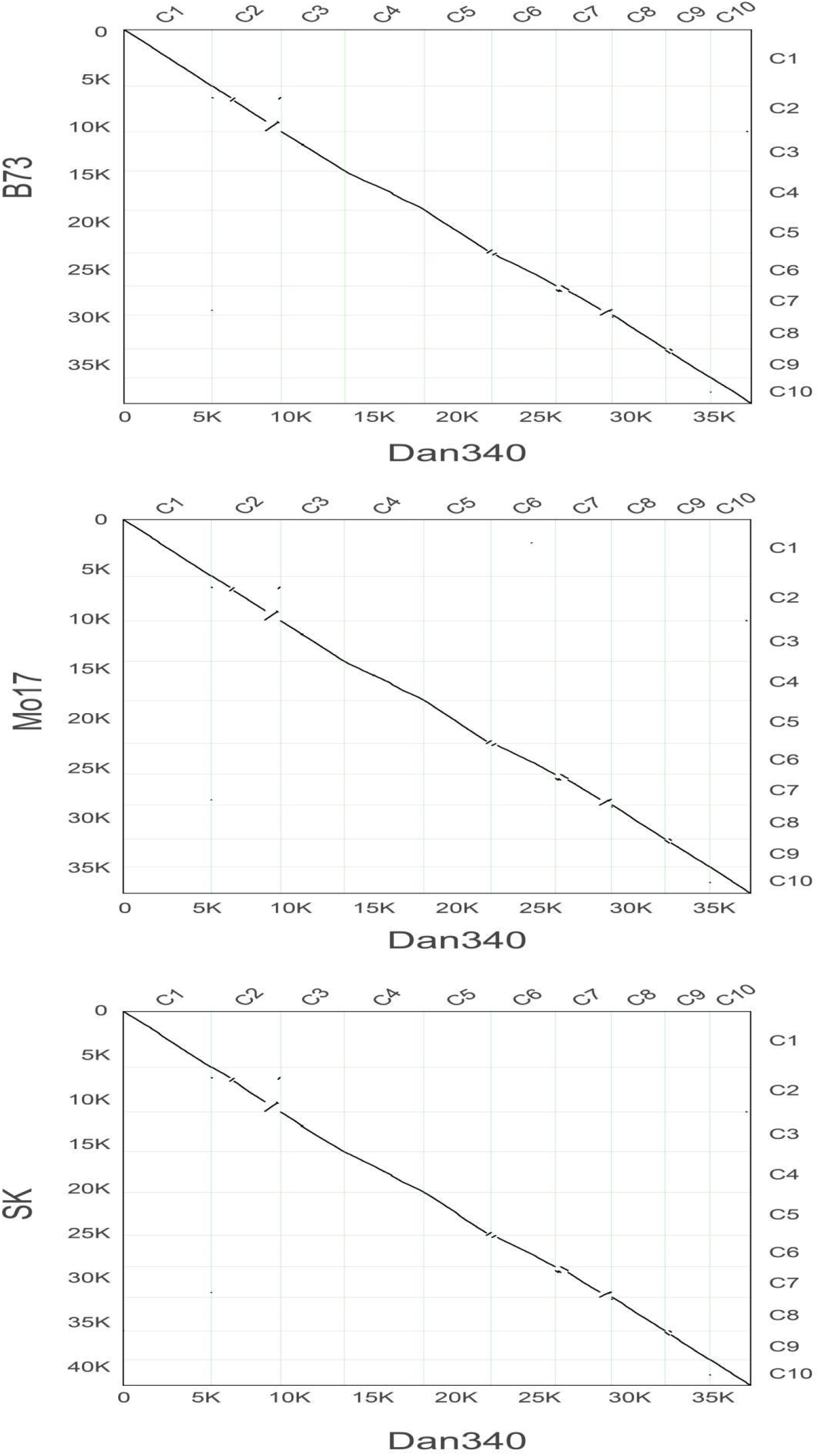
Pairwise comparison of genome sequences using a dot plot between the Dan340 line and B73 (23,350 gene pairs), Mo17 (21,913 gene pairs), SK (23,016 gene pairs). The horizontal axis represents the target species, the vertical axis represents the reference species, C1 – C10 represent the respective chromosomes 1 – 10, 0 – 35 k represent the chromosome length scale marks, which mainly reflect the lengths of the chromosomes, and a point represents a pair of common genes.

**Table 2.** LAI scores of the four tested maize inbred lines.

| Lines | ~~D340~~-Dan340 | B73 | Mo17 | SK |
| --- | --- | --- | --- | --- |
| LAI | 25.13 | 24.94 | 24.45 | 27.12 |

### Genome annotation

Repeat sequences of the Dan340 genome were annotated using both *ab initio* and homolog-based search methods. For the *ab initio* prediction, RepeatModeler v1.0.8 [33] (RRID:SCR_015027), RepeatScout version 1.0.5 [34] (RRID:SCR_014653), and LTR_FINDER version 1.07 [31] were used to discover transposable elements (TEs) and to build TEs library. An integrated TEs library and a known repeat library (Repbase V15.02, homolog-based)(RRID:SCR_021169) were subjected to RepeatMasker v3.3.0 [35] (RRID:SCR_012954) to predict TEs. For homolog-based predictions, RepeatProteinMask was performed to detect TEs in the genome by comparing it against a TE protein database. Tandem repeats were ascertained in the genome using Tandem Repeats Finder (TRF, version 4.07b) [36] (RRID:SCR_022193). As a result, 1723.99 Mb of repeat sequences were identified, accounting 73.40% of the genome size. Among these repeat sequences, 1,555.57 Mb were predictedas to be long-terminal repeat (LTR) retrotransposons and 44.53 Mb were predictedas to be DNA transposons, accounting for 66.23% and 1.60% of the genome, respectively. Furthermore, among the LTR retrotransposons, the Gypsy and Copia superfamilies comprised 23.81% and 12.75% of the genome, respectively. Thus, retrotransposons accounted for a large proportion of the Dan340 genome, which was consistent with the genomic characteristics of other maize inbred lines (Table 2). All repetitive regions except tandem repeats were soft-masked for protein-coding gene annotation. Five *ab initio* gene prediction programs, Augustus v3.0.2 [37–39] (RRID:SCR_008417), Genscan v1.0 [40] (RRID:SCR_013362), Geneid v1.4 [28], GlimmerHMM v3.0.2 [41] (RRID:SCR_002654) and SNAP v2013-02-16 [42] (RRID:SCR_007936), were used to predict genes. In addition, the protein sequences of five homologous species (*Sorghum bicolor, Setaria italica, Hordeum vulgare, Triticum aestivum*, and *Oryza sativa*) were downloaded from Ensembl and NCBI. Homologous sequences were aligned against the genome using TBLASTN (E-value 1E-05). Genewise version 2.2.0 [27] was employed to predict gene models on the basis of the sequence alignment results.

For RNA-seq prediction, fresh samples of six tissues (stem, endosperm, embryo, bract, silk, and ear tip) were collected. Total RNA was extracted from each sample using an RNAprep Pure Plant Kit (Tiangen Biotech Co., Ltd., Beijing, China). Isolated purified RNA was the template for the construction of a cDNA library, having fragment lengths of approximately 300 bp, using the NEBNext Ultra RNA Library Prep Kit for Illumina (New England Biolabs, Ipswich, MA, USA) in accordance with the manufacturer’s instructions. Sequencing was performed on an Illumina HiSeq X Ten platform and 150-bp paired-end reads were generated. Raw reads were trimmed by removing adapter sequences, reads with more than 5% of unknown base calls (N), and low-quality bases (base quality less than 5, Q ≤5). Clean paired-end reads were aligned to the genome using Tophat version 2.0.13 [43] (RRID:SCR_013035) to identify exon regions and splice positions. The alignment results were then used as input for cufflinks version 2.1.1 [44] (RRID:SCR_014597) to assemble transcripts to the gene models. In addition, RNA-seq data were assembled using Trinity version 2.1.1 [45] (RRID:SCR_013048), creating several pseudo-ESTs. These pseudo-ESTs were also mapped to the assembled genome using BLAT and gene models were predicted using PASA [46] (RRID:SCR_014656). A weighted and non-redundant gene set was generated using EVidenceModeler (EVM, version 1.1.1) [47] (RRID:SCR_014659), which merged all the gene models predicted by the above three approaches. Finally, PASA was used to adjust the gene models generated by EVM. As a result, a total of 39,733 protein-coding genes were annotated in the final set. To better understand gene functions, we used all 39,733 protein-coding genes as query against public protein databases, including NCBI non-redundant protein sequences (Nr), Swiss-Prot, Protein family (Pfam), Kyoto Encyclopedia of Genes and Genomes (KEGG), InterPro, and Gene Ontology (GO). In total, 39,646 genes (99.8%) could be annotated from these databases and 24,402 genes (61.41%) were supported by RNA-seq data. Furthermore, the gene number, gene length distribution, and exon length distribution were all comparable to those of other maize inbred lines and common crop species (Table 3).

**Table 3.** Summary statistics of annotated protein-coding genes in Dan340 and other maize inbred lines and common crop species.

| Species | Number | Average | Average | Average | Average | Average |
| --- | --- | --- | --- | --- | --- | --- |
|  |  | transcript | CDS | exons per | exon | intron |
|  |  | length(bp) | length(bp) | gene | length(bp) | length(bp) |
| Dan340 | 39,733 | 3,793.47 | 1,140.91 | 4.69 | 243.47 | 719.61 |
| B73 | 39756 | 3511.78 | 1102.11 | 4.58 | 240.64 | 673.10 |
| Mo17 | 38620 | 3362.68 | 1140.26 | 4.69 | 242.98 | 601.83 |
| SK | 42942 | 3857.18 | 1179.17 | 4.83 | 243.93 | 698.48 |
| Hvu | 24,286 | 2,116.13 | 1,093.77 | 4.1 | 267.02 | 330.19 |
| Osa | 35,679 | 2,165.58 | 991.55 | 3.78 | 262.57 | 422.87 |
| Sbi | 34,008 | 2,626.44 | 1,164.14 | 4.31 | 270.09 | 441.76 |
| Sit | 27,233 | 2,982.22 | 1,336.29 | 5.14 | 260.2 | 397.98 |
| Tae | 103,539 | 3,087.61 | 1,277.31 | 4.51 | 283.23 | 515.78 |

Abbreviations: Hvu: *Hordeum vulgare*; Osa: *Oryza sativa*; Sbi: *Sorghum bicolor*; Sit: *Setaria italica*; Tae: *Triticum aestivum*.

Transfer RNA (tRNA) genes were predicted using tRNAscan-SE software v. 1.4 [48] (RRID:SCR_010835) with the default parameters. Ribosomal RNAs (rRNAs) were annotated on the basis of their homology levels with the rRNAs of several species of higher plants using BLASTN with an E-value of 1e-5. The microRNA (miRNA) and small nuclear RNA (snRNA) fragments were identified by searching the Rfam database v. 11.0 (RRID:SCR_007891) using INFERNAL v. 1.1 software (RRID:SCR_011809) [49, 50]. Finally, 4,547 miRNAs, 5,963 tRNAs, 63,564 rRNAs, and 1,422 snRNAs were identified, which had average lengths of 126.79, 75.25, 309.47, and 132.10 bp, respectively (Table 4).

**Table 4.**
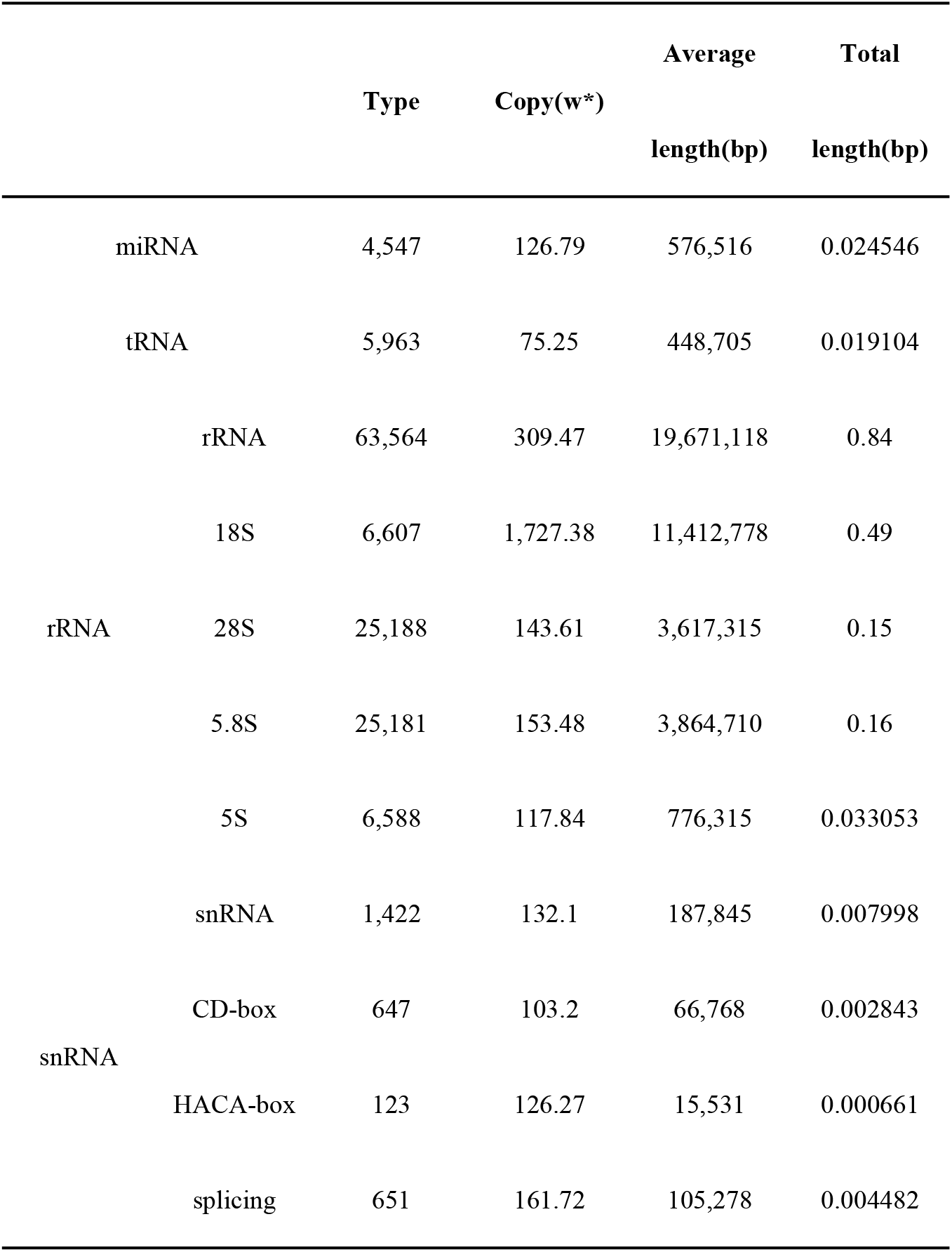
Annotation statistics of non-coding RNAs in the Dan340 genome using different databases.

### Comparative genomic analysis between Dan340 and other maize lines

We applied the OrthoMCL pipeline [51] to identify orthologous gene families among the four maize inbred lines, including Dan340, B73, Mo17, and SK. The longest protein from each gene was selected, and the proteins with a length less than 30 amino acids were removed. Subsequently, pairwise sequence similarities between all input protein sequences were calculated using BLASTP with an E value cut-off of 1×10^-5^. Markov clustering (MCL) of the resulting similarity matrix was used to define the ortholog cluster structure of the proteins, using an inflation value (-I) of 1.5 (default setting in OrthoMCL). Next, comparative analyses were performed among Dan340, B73, Mo17, and SK (Fig. 6A).

**Figure 6.**
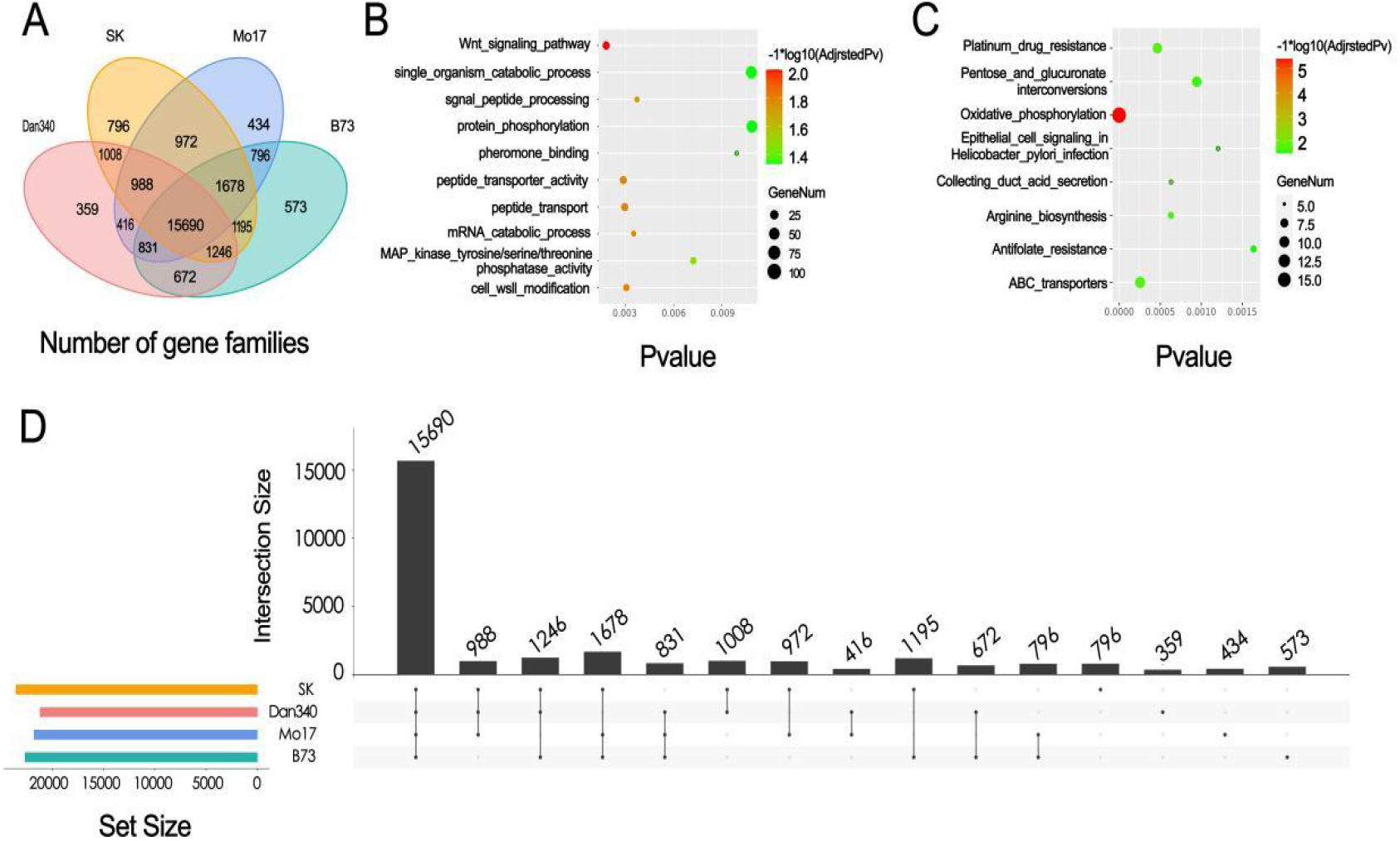
Gene family analyses and core- and pan-genomes of maize. A, Comparisons of gene families in Dan340, B73, Mo17, and SK. The Venn diagram illustrates shared and unique gene families among the four maize inbred lines. B, Gene ontology enrichment analysis of Dan340-specific genes. C, Kyoto Encyclopedia of Genes and Genomes analysis of Dan340-specific genes. D, Core- and pan-genome of maize. The histograms show the core-gene clusters (shared in all four genomes), dispensable gene clusters (present in three or two genomes) and specific gene clusters (present only in one genome).

Analysis of genes in the Dan340 genome, together with those from B73, Mo17 and SK, were clustered into 27,654 gene families. Of these, 15,690 families were shared among the four maize inbred lines, representing a core set of genes across these maize genomes. There were 1,806 genes from 359 gene families that were specific to Dan340, of which many had functional gene ontology annotations relating to “protein phosphorylation”, “single-organism catabolic process”, and “pheromone binding” (Fig. 6B). In KEGG functional enrichment, the most enriched pathway of Dan340 specific genes were “antifolate resistance”, “epithelial cell signaling in helicobacter pylori infection”, and “pentose and glucuronate interconversions” (Fig. 6C).

In addition, OrthoMCL was used to identify the core and dispensable gene sets on the basis of gene family. The gene families that were shared among the four inbred lines were defined as core gene families. Furthermore, gene families that were shared among three inbred lines, between two inbred lines, and those that were only present in one inbred line (private gene families) were also displayed in Fig 6D.

### Genetic variation analysis

To investigate the genetic variations between Dan340 and other maize inbred lines, we used PBSV version 2.2.2 [52] to detect structural variations. First, PacBio reads of B73 and Mo17 were downloaded from MaizeGDB [53], and PacBio reads of SK were obtained from the National Genomics Data Center [54]. Next, subreads were aligned to the Dan340 reference genome assembled in this study using pbmm2 to generate a bam file. Then, Samtools v1.7 [55] (RRID:SCR_002105) was used to identify and split the bam file on the basis of chromosomes and scaffolds. Afterwards, pbsv discover was used to generate svsig result files for different chromosomes and scaffolds. The svsig files of multiple samples were then used together to perform the SV joint calling with “pbsv call”, and finally, the vcf files were obtained.

The high-quality Dan340 reference genome allowed us to identify large SVs presented in different maize inbred lines. By mapping the PacBio long-reads of B73 to Dan340 genome, we identified a total of 8,289 structural variations (length longer than 500 bp) between the two representative maize genomes, including 1,653 insertions, 6,537 deletions, 36 inversions, and 63 duplications (Table 5). Furthermore, the structural variations presented in Mo17 and SK were also detected in this study (Table 6 & 7). This dataset provides abundant variation resources for molecular improvement and breeding in maize in the future.

**Table 5.** Structural variations between Dan340 and B73.

| Chr.<br>Number | Insertion |  | Deletion |  | Inversion |  | Duplication |  |
| --- | --- | --- | --- | --- | --- | --- | --- | --- |
|  | Number | Length<br>(bp) | Number | Length<br>(bp) | Number | Length<br>(bp) | Number | Length<br>(bp) |
| 1 | 266 | 432723 | 998 | 13194452 | 7 | 120136 | 8 | 311433 |
| 2 | 183 | 316530 | 743 | 10075332 | 7 | 177706 | 9 | 261774 |
| 3 | 201 | 381536 | 838 | 11475573 | 5 | 24457 | 7 | 37754 |
| 4 | 183 | 303142 | 698 | 9809218 | 2 | 113807 | 7 | 255958 |
| 5 | 181 | 330690 | 651 | 9519227 | 3 | 108429 | 6 | 121490 |
| 6 | 122 | 252117 | 527 | 7723482 | 3 | 55199 | 8 | 300610 |
| 7 | 140 | 226942 | 512 | 7146020 | 2 | 49011 | 4 | 122886 |
| 8 | 140 | 204697 | 577 | 7720113 | 1 | 80400 | 3 | 42148 |
| 9 | 121 | 219627 | 569 | 7539785 | 3 | 17955 | 9 | 114595 |
| 10 | 116 | 209091 | 424 | 6256310 | 3 | 130724 | 2 | 40306 |

**Table 6.** Structural variations between Dan340 and Mo17.

| Chr. | Insertion |  | Deletion |  | Inversion |  | Duplication |  |
| --- | --- | --- | --- | --- | --- | --- | --- | --- |
|  | Number | Length (bp) | Number | Length (bp) | Number | Length (bp) | Number | Length (bp) |
| 1 | 171 | 226204 | 1049 | 14591535 | 10 | 223256 | 6 | 157653 |
| 2 | 102 | 164536 | 668 | 9437198 | 4 | 40341 | 5 | 149456 |
| 3 | 121 | 196809 | 767 | 11363033 | 0 | 0 | 4 | 26512 |
| 4 | 96 | 146230 | 628 | 8996741 | 3 | 50713 | 10 | 169360 |
| 5 | 102 | 147883 | 660 | 9449523 | 3 | 190422 | 7 | 85046 |
| 6 | 75 | 101725 | 567 | 8423089 | 3 | 130693 | 3 | 20573 |
| 7 | 84 | 119614 | 521 | 7784957 | 4 | 159203 | 6 | 275006 |
| 8 | 83 | 123397 | 603 | 8751739 | 2 | 30781 | 5 | 214099 |
| 9 | 61 | 83589 | 529 | 7194721 | 1 | 10328 | 3 | 55228 |
| 10 | 75 | 126350 | 497 | 6832836 | 5 | 227243 | 4 | 159642 |

**Table 7.** Structural variations between Dan340 and SK.

| Chr. | Insertion |  | Deletion |  | Inversion |  | Duplication |  |
| --- | --- | --- | --- | --- | --- | --- | --- | --- |
|  | Number | Length (bp) | Number | Length (bp) | Number | Length (bp) | Number | Length (bp) |
| 1 | 244 | 355348 | 1226 | 17656467 | 15 | 228724 | 16 | 404487 |
| 2 | 133 | 220966 | 842 | 12542190 | 6 | 244805 | 5 | 76672 |
| 3 | 173 | 286569 | 894 | 12961969 | 4 | 7657 | 6 | 96601 |
| 4 | 211 | 338458 | 1012 | 14231859 | 3 | 70762 | 14 | 315182 |
| 5 | 160 | 248233 | 786 | 10660145 | 8 | 280670 | 5 | 206286 |
| 6 | 93 | 118612 | 588 | 8923936 | 4 | 55606 | 2 | 1760 |
| 7 | 115 | 175971 | 622 | 8865791 | 1 | 30856 | 9 | 233675 |
| 8 | 143 | 210185 | 685 | 10443437 | 2 | 3061 | 6 | 152412 |
| 9 | 114 | 186005 | 647 | 8682441 | 4 | 9595 | 7 | 37664 |
| 10 | 109 | 159668 | 559 | 8726188 | 4 | 132337 | 4 | 95237 |

## Conclusions

We assembled the chromosome-level genome of the maize elite inbred line Dan340 using long CCS reads from the third-generation PacBio Sequel II sequencing platform, with scaffolding informed by chromosomal conformation capture (Hi-C). The final assembly of the Dan340 genome was 2,348.72 Mb, including 2,738 contigs and 2,315 scaffolds with N50 of 41.49 Mb and 215.35 Mb, respectively. Comparisons of the Dan340 genome with the reference genomes of three other common maize inbred lines identified 1,806 genes from 359 gene families that were specific to Dan340. In addition, we also obtained large numbers of structural variants between Dan340 and other maize inbred lines, and these may be underlying the mechanisms responsible for the phenotypic discrepancies between Dan340 and other maize varieties. Therefore, the assembly and annotation of this genome not only facilitates our understanding of the intraspecific genomic diversity in maize, but they also serve as novel resources for maize breeding improvement.

## Data Availability

The raw sequence data have been deposited in NCBI under project accession No. PRJNA795201. Data is also available in the *GigaScience* GigaDB repository [56].

## Abbreviations

BUSCO: Benchmarking Universal Single-Copy Orthologs
Hi-C: chromosomal conformation capture
CCS: circular consensus sequencing
HiFi: long high-fidelity
CEGMA: Core Eukaryotic Genes Mapping Approach
LTR: long-terminal repeat
LAI: long-terminal repeat assembly index
TEs: transposable elements
TRF: Tandem Repeats Finder
EVM: EVidenceModeler
KEGG: Kyoto Encyclopedia of Genes and Genomes
GO: Gene Ontology
MCL: Markov clustering
NCBI: National Center for Biotechnology Information
PacBio: Pacific Biosciences
tRNA: Transfer RNA
rRNAs: Ribosomal RNAs
miRNA: microRNA
snRNA: small nuclear RNA

## Authors’ contributions

F.W., J.Z. and H.Z. conceived the project; Y-K.Z. D.M. and Y.W. wrote, modified the manuscript; L.X. G.F. and L.W. performed the data curation; Y.H. L.Z. Y-L.Z. and Z.L. analyzed data. All authors read and approved the final manuscript.

## Acknowledgements

This research was supported by grants from the special project for the construction of scientific and technological innovation capacity of Beijing Academy of Agriculture and Forestry Sciences (NO. KJCX20200305).

